# HairSplitter: haplotype assembly from long, noisy reads

**DOI:** 10.1101/2024.02.13.580067

**Authors:** Roland Faure, Dominique Lavenier, Jean-François Flot

## Abstract

**Motivation:** Long-read assemblers face challenges in discerning closely related viral or bacterial strains, often collapsing similar strains into a single sequence. This limitation has been hampering metagenome analysis, as diverse strains may harbor crucial functional distinctions.

**Results:** We introduce a novel software, HairSplitter, designed to retrieve strains from a partially or totally collapsed assembly and long reads. The method uses a custom variant-calling process to operate with erroneous long reads and introduces a new read binning algorithm to recover an a priori unknown number of strains. On noisy long reads, HairSplitter recovers more strains while being faster than state-of-the-art tools, both in the cases of viruses and bacteria.

**Availability:** HairSplitter is freely available on GitHub at github.com/RolandFaure/HairSplitter.

## Introduction

Microbiomes play a crucial roles in many ecosystems, such as soils or human guts, in turn impacting human health (Conlon and Bird, 2014) and soil fertility (Coban et al., 2022). Microbiomes typically contain sets of organisms with highly similar genomes, the sequences of which are called haplotypes (short for “haploid genotypes” (Ceppellini et al., 1967)). Distinguishing these lineages is an important challenge, as small genomic differences between haplotypes can lead to significant phenotypic changes. For instance, some strains of *Escherichia coli* can be pathogenic or commensal while having an Average Nucleotide Identity (ANI) (Konstantinidis and Tiedje, 2005) of more than 98.5% (Frank et al., 2011). A few mutations also became famous for altering significantly the infectiousness of some coronaviruses lineages (Magazine et al., 2022).

*De novo* sequencing and assembling is a central method to characterize microbial communities. Unlike previous methods, it allows to analyse the composition of a metagenome without culturing the strains, enabling a wide range of analyses (Ward, 2006). While existing genome assemblers proficiently reconstruct genomes of abundant species, they struggle to distinguish viral or bacterial haplotypes. The main difficulty for assemblers lies in the unknown number of haplotypes in a sample and their uneven coverage (Ghurye et al., 2016).

Many tools have been developed to overcome this problem in the context of short-read assemblies, such as OPERA-MS (Bertrand et al., 2019), Constrains (C Luo et al., 2015), STRONG (Quince et al., 2021), StrainXpress (Kang et al., 2022) and VStrains (R Luo and Lin, 2023). However, these methods are not designed for long-read sequencing and do not exploit the long-range information contained in long reads.

Long reads with extremely low error rate, such as PacBio HiFi reads, have been used to distinguish finely strains with the help of specialized software such as hifiasm (Cheng et al., 2021) and stRainy (Kazantseva et al., 2023). However, this challenge has not been yet successfully tackled in the case of noisier reads such as “regular” PacBio data or Oxford Nanopore Technology (ONT) reads, the latter of which can be obtained very rapidly on cheap sequencers that are small enough to be carried into the field (Cesare et al., 2024; Runtuwene et al., 2019).

Several methods have been implemented to deal with haplotype reconstruction from long reads with high error rates. While the viral and bacterial haplotype assembly problems are identical in their formulation, the characteristics of the input data vary significantly: the genomes are generally much shorter and much more deeply sequenced in the case of viruses. This has led to the emergence of software specialized in either one of the two problems. In the context of bacterial strain separation, Vicedomini et al., 2021 showed that mainstream assemblers such as metaFlye (Kolmogorov et al., 2020) and Canu (Koren et al., 2017) failed to distinguish close bacterial haplotypes and proposed a new tool, called Strainberry, to reconstruct strains. In the context of viral strain separation, Strainline (X Luo et al., 2022) and HaploDMF (Cai et al., 2022) were presented to tackle specifically the viral haplotype reconstruction problem and need very high depth of sequencing to work. The method iGDA (Z Feng et al., 2021) was proposed as a general approach to phase minor variants while handling high error rates and can theoretically assemble both bacterial and viral haplotypes. The main shortcomings of all of these methods is that they struggle to recover haplotypes of low abundance. Additionally, most of these tools are very computationally intensive.

We present HairSplitter, an efficient pipeline for separating haplotypes in viral and bacterial context using error-prone long reads. HairSplitter first calls variants using a custom process to distinguish actual variants from alignment or sequencing artefacts ; clusters the reads into an unspecified number of haplotypes ; creates the new separated contigs ; and finally untangles the assembly. HairSplitter can be used for either metaviromes or bacterial metagenomes.

## Methods

### Overview of the pipeline

HairSplitter takes as input an assembly (in fasta format) or an assembly graph (in gfa format) as well as sequencing reads (fasta/q) and produces a new assembly (fasta and gfa). The HairSplitter pipeline is depicted in Figure 1 and comprises five steps: 1) correcting the assembly, 2) calling variants on each contig, 3) clustering the reads by haplotype on each contig, 4) reassembling the strain-specific contigs and 5) unzipping.

**Figure 1.**
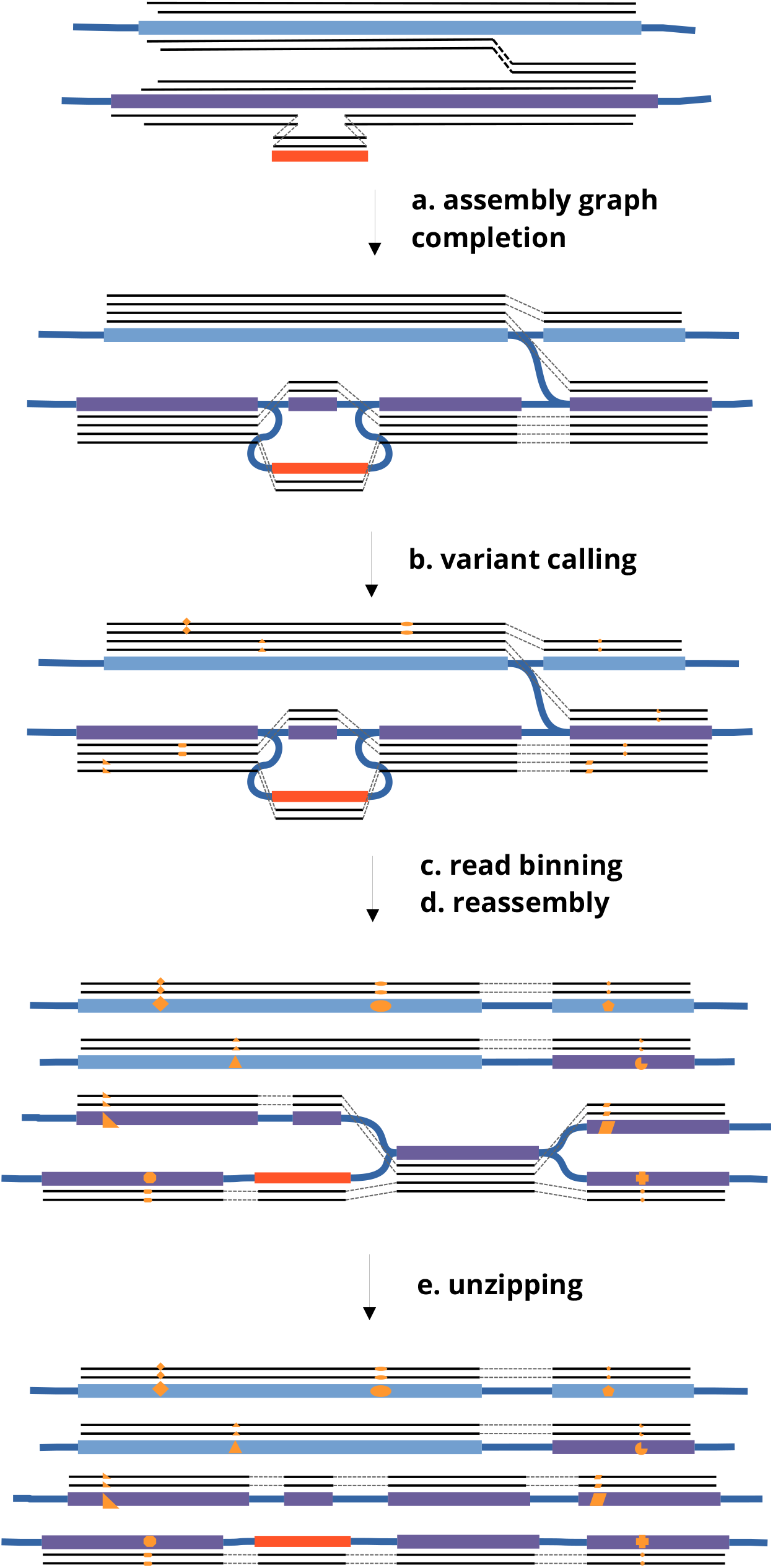
Illustration of the five steps of the HairSplitter pipeline. Colored rectangles represent contigs, thick blue lines are links in the assembly graph and black lines represent the reads aligned on the assembly. Orange shapes on reads and contigs indicate variant positions compared to the original sequence.

### Completion of the assembly graph

To work well, HairSplitter needs as input an assembly graph on which all genomic reads align from end to end, which we define as a “complete” assembly graph. If the assembly was not provided as a graph, it is turned into a graph with no edges. Collapsed assembly graphs are often incomplete because of contigs that have been detached from their neighbors and of collapsed structural variation between strains.

Aligning reads on an incomplete graph reveals locations where a significant number of reads stop aligning, which we call breakpoints. Breakpoints can occur in the middle or the end of contigs. To complete the initial assembly graph, the reads are aligned on the graph using minigraph (Li et al., 2020). The assembly is subsequently examined for breakpoints and HairSplitter breaks the contigs at these breakpoints. Additionally, links are added in the graph between ends of contigs when there is sufficient read support. The process is illustrated in Figure 1a. An evaluation of this step in terms of misassemblies and contiguity is provided in Supplementary Table 4.

The completed assembly resulting from this process is used throughout the subsequent stages of the pipeline.

### Mathematical model behind variant calling

To sort reads into haplotypes, the intuitive method of clustering reads based on the similarity of their full sequence proves ineffective due to the prevalence of sequencing and alignment errors, obscuring strain differences. HairSplitter first identifies variant positions, pinpointing loci where strains exhibit actual differences. The reads are then separated based only on these loci. We did not find any variant caller suitable for our specific challenge - calling variants with noisy long reads in a metagenomic context including potentially low-abundance strains while maintaining high computational efficiency. Thus, we devised our own variant calling procedure.

The naivest procedure to identify polymorphic loci consists in going through the pileup of the reads on the assembly and identifying loci where at least a proportion *λ* of reads have an alternative allele. However, this approach falls short when using error-prone reads. For instance, in the case of a strain representing only 1% of the total of the reads, *λ* needs to be less than 0.01 to detect variant positions corresponding to this strain, resulting in the selection of many artefactual positions if the reads have an error rate *>* 1%.

The key lies in taking several loci into account simultaneously, an idea already explored in Z Feng et al., 2021 and leveraging the assumption that alignment artifacts occur randomly in the pileup while genomic variant are expected to be correlated along the alignment. Consequently, pileups at polymorphic loci are expected to exhibit strong correlation, contrary to pileups at non-polymorphic loci. HairSplitter introduces a new statistical approach and a new algorithm to exploit this observation and detect even rare strains, as illustrated below.

Consider a complete pileup of *n* reads over *m* positions, which we will model as a matrix of letters. Let us assume that errors occur independently on all reads and at all positions with a probability *≤ ϵ* and that all errors in a given column are identical (worst-case scenario). We aim to estimate the probability that there exist *a* reads that share errors at *b* different loci: in other words, the probability that there exist a submatrix of size *a *b* containing only errors in the pileup, defined by selecting *a* rows (reads) and *b* columns (loci).

There exist 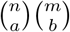 submatrices of size *a ∗ b*. Each of these submatrix has a probability lower than 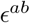 to contain only errors. Therefore, given that expectation is linear (DeGroot and Schervish, 2002), the expectation *E* of the number of submatrices of size *a ∗ b* containing only errors in the pileup is lower than 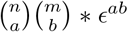. Now, to obtain the probability that there exist no submatrix of size *a ∗ b* containing only errors, we can use Markov’s inequality, according to which the probability that a positive random variable be higher than 1 is always smaller than the expectation of this variable (DeGroot and Schervish, 2002). Here, it tells us that the probability that there exist a submatrix containing only errors is smaller than *E*. In other terms, the probability that there exist somewhere in the pileup *a* reads sharing errors at *b* different loci is lower than 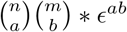.

Now, let us consider a pileup with *n* = 1000 reads across *m* = 5000 positions and *ϵ* = 0.1. The probability that there exist *a* = 10 reads sharing errors at *b* = 10 different loci is lower than 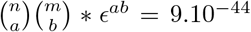. Therefore, if the error rate is of 10% or less and the pileup indicates 10 reads (1% coverage) sharing an alternative allele at 10 loci (divergence of 0.2%), we can confidently assume that these are not errors, suggesting these reads originate from the same strain, and the loci are polymorphic sites.

Despite its simplistic nature, this model underscores the statistical power gained by examining multiple loci simultaneously, enabling the detection of low-abundance, highly similar strains even in the presence of very noisy long reads. The idea behind the model is illustrated in Figure 2.

**Figure 2.**
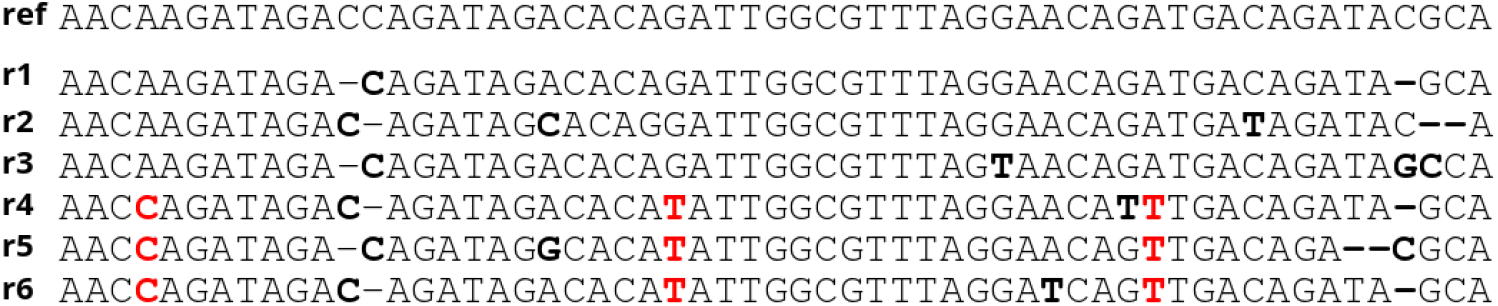
In this pileup of reads, does the submatrix of variants highlighted in red indicate the presence of two strains? The probability that there exist 3 reads having an alternative allele at 3 loci if we estimate *e* = 0.1 is less than 0.02: the variants are thus likely not independent and probably underline the presence of at least two different strains.

### Variant calling

The approach to identifying polymorphic loci capitalizes on the statistical power underlined above. Specifically, HairSplitter aims to identify clusters of positions featuring alternative alleles on the same reads.

To generate the pileup, all reads are aligned to the contigs of the completed assembly using minimap2 (Li, 2018). HairSplitter then traverses the pileup of each contig and determines, for each position, the majority allele and the main alternative allele (either a base or an indel). Long indels are treated as multiple adjacent loci. Only positions with a minimum of five reads carrying alternative alleles are considered as potential polymorphic sites to ensure statistical robustness (cf. model above). HairSplitter compares each new position to previously observed positions. If the set of reads with alternative alleles at this position and at a previously encountered position share more than 90% reads, the new position is clustered with the old one.

After all positions have been considered, clusters are tested using the statistical model described above and only clusters with a p-value below 0.001 are kept. The corresponding positions are outputted as polymorphic sites.

### Read binning

The contig is divided into windows with a default size of *w* (by default, 2000 bases). Reads are binned into haplotypes sequentially on the windows of a contig. Only reads spanning the entirety of the window are considered for binning. To cluster reads, HairSplitter operates on the premise that reads originating from the same haplotype should be identical at all polymorphic loci. Nevertheless, inherent sequencing and variant-calling errors might introduce unintended discrepancies among reads from a single haplotype. To address this, HairSplitter adopts a three-step strategy.

Step one is to correct errors at polymorphic loci. HairSplitter corrects the errors at polymorphic loci by performing a k-nearest-neighbour imputation (Fix and Hodges, 1989), with *k* = 5. The distance between two reads is defined as the number of different alleles at polymorphic positions. Each base of the pileup is considered and changed to the most frequent base among the *k* nearest neighbours on all reads and all positions until convergence.

Step two is to form clusters of reads, clustering reads together if and only if they exhibit no differences at any polymorphic loci. Indeed, two reads that bear at least one different allele originate by definition from two different haplotypes.

In the third step, a last check is run to rescue small clusters that can arise from errors in Step 1. HairSplitter constructs a graph linking each read to its *k* closest neighbours, including links between all pairs of reads differing on one position or less. The graph is then clustered using the Chinese Whispers algorithm (Biemann, 2006), initialising the clustering with the clusters obtained in the second step. The Chinese Whispers algorithm iteratively assign reads to the most represented cluster among their neighbors until convergence. The Chinese Whispers algorithm always converge toward a stable solution, i.e. a clustering where all reads are in the same group as at least half of their neighbors. There exist many stable clusterings but the algorithm is likely to converge to a solution close to the initialization: the clusters obtained in the second step are unlikely to be significantly altered, but very small clusters will likely be merged with other close clusters.

### Reassembly

Across all windows on every contig, the original sequence undergoes repolishing using the haplotype-specific groups of reads previously identified. The polishing can be executed with either Racon (Vaser et al., 2017) or Medaka (github.com/nanoporetech/medaka), with the latter being more precise but considerably slower in our experience. By default, HairSplitter uses Medaka only for short genomes (*≤* 1 Mb).

### Graph Unzipping

The resulting assembly comprises contigs of length *w* that can easily be stitched into longer contigs. For this purpose, a straightforward algorithm is employed, GraphUnzip (Faure et al., 2021), depicted in Figure 1e. Let us call a contig exhibiting multiple outgoing links with other contigs at one end a “knot”. Knots generally represent collapsed contigs. GraphUnzip initially aligns all reads on the assembly graph. Subsequently, GraphUnzip iteratively assess nodes. If more than three reads traverse a neighbor of the knot (called A), then traverse the knot, and traverse another neighbor at the opposite end of the knot (called B), the knot is duplicated to create a new contig that has as unique neighbors A and B. The links from A and B to the original knot are deleted, preserving only the links to the copy of the contig. This process is repeated until no further knots can be duplicated.

## Results

### Datasets

The datasets used in this article are described in Table 1. The accession numbers of the data in public repositories can be found in section “Reproducibility and data availability”.

**Table 1.** Characteristics of the different datasets used for benchmarking on real data.

| dataset | species | # strains | strain coverages | ANI divergence | sequencing technology |
| --- | --- | --- | --- | --- | --- |
| HBV-2 | Hepatitis B | 2 | 4000x, 9900x | 10% | Nanopore R.9.4.1 |
| norovirus-7 | norovirus | 7 | 50, 350, 450, 700, 900, 1150, 1400x | 1-3.9 % | Nanopore R.9.4.1 |
| *V. fluvialis* | *Vagococcus fluvialis* | 5 | 90x, 136x, 172x, 182x, 206x | 0.01-1.51% | Nanopore R9.4.1 |
| Zymo-GMS Q9 | *Escherichia coli* | 5 | 90x, 90x, 90x, 90x, 90x | 0.37-1.51% | Nanopore R9.4.1 |
| Zymo-GMS Q20 | *Escherichia coli* | 5 | 25x, 25x, 25x, 25x, 25x | 0.37-1.51% | Nanopore R10.4.1 |
| Zymo-GMS HiFi | *Escherichia coli* | 5 | 41x, 41x, 41x, 41x, 41x | 0.37-1.51% | PacBio HiFi |

### Bacterial datasets

We used the Zymobiotics Gut Microbiome Standard (abbreviated to Zymo-GMS) and a *Vagococcus fluvialis* dataset (Rodriguez Jimenez et al., 2022) to compare the performance of different algorithms designed to separate bacterial haplotypes in a metagenomic context. Zymo-GMS is a mixture of bacteria, archaea and yeast (21 different strains in total) dosed to mimic the composition of the human gut microbiome. These 21 strains include five *Escherichia coli* strains, which we used to evaluate the strain-separation ability of various programs. Three Zymo-GMS sequencing were used, respectively from a Nanopore R9.4.1 run, a Nanopore 10.4.1 run and a PacBio HiFi run. The *Vagococcus fluvialis* dataset consists of a mix of five *Vagococcus fluvialis* strains that were sequenced together using barcoded reads, each barcode corresponding to a strain. We did not use the barcode information for the assemblies, reserving them for validation. Among the five strains, three had an Average Nucleotide Idendity (ANI) over 99.99%. metaFlye is used to assemble the reads, as it yielded better assemblies compared to Canu according to Vicedomini et al. (Vicedomini et al., 2021).

In addition, we simulated datasets to assess the impact of the number of strains, coverage and divergence on the assemblies. These experiments were directly inspired by the protocol of Vicedomini et al. (Vicedomini et al., 2021). The genomes of ten strains of *Escherichia coli* were downloaded from the SRA, namely 12009 (GCA_000010745.1), IAI1 (GCA_000026265.1), F11 (GCA_018734065.1), S88 (GCA_000026285.2), Sakai (GCA_ 003028755.1), SE15 (GCA_000010485.1), *Shigella flexneri* (GCF_000006925.2), UMN026 (GCA_000026325.2), HS (GCA_ 000017765.1), and K12 (GCF_009832885.1). These strains were chosen to be representative of the diversity of *E. coli*. We simulated Nanopore sequencing using Badread (R Wick, 2019) with the setting “Nanopore2023” to simulate 50x of R10.4.1 reads. Between 2 and 10 strains were mixed to assess how many strains the software could separate. From the 10-strain mix, the 12009 strain was downsampled to 30x, 20x, 10x and 5x to assess the impact of the coverage on strain separation. Finally, to assess the impact of the divergence of sequences on strain separation, 50x of reads were simulated for strain K12 and for strains of decreasing divergence with K12; assemblies of reads of K12 mixed with reads of each of these strain were evaluated for separation.

### Viral datasets

Two datasets were used to benchmark the performance of the programs tested at separating viral haplotypes, a ix of two strains of hepatitis B Virus (HBV) from McNaughton et al., 2019 and an in-silico mix of the sequencing of seven strains of norovirus from Flint et al., 2021. These datasets were directly taken from the paper of HaploDMF (Cai et al., 2022). The reference genomes to run reference-based tools were taken as the reference genome in the GenBank database, GCF_000861825.2 for HBV and MW661279.1 for norovirus.

### Performance evaluation

We used MetaQUAST (Mikheenko et al., 2016) to measure assembly features such as assembly length, NG50, misassemblies, mismatches, indels and completeness. MetaQUAST was run with the –unique-mapping and –reuse-combined-alignments options to prevent a sequence, whether a contig or part of it, from being mapped to multiple distinct reference locations.

To assess if strains are well separated, the most important metric is the completeness of the resulting assembly. We chose to assess MetaQUAST completeness but also 27-mer completeness. MetaQUAST completeness measures the percentage of the solution on which the assembly aligns, while 27-mer completeness measures the percentage of the 27-mers of the solution that are effectively found in the assembly. Collapsed homozygous contigs typically impact negatively MetaQUAST completeness but not 27-mer completeness.

### Evaluated software

In addition of HairSplitter, we chose to evaluate the programs stRainy (Kazantseva et al., 2023) and Strainberry (Vicedomini et al., 2021), which have been introduced specifically as bacterial strain separation methods; hifiasm-meta (X Feng et al., 2022), which is the most popular assembler for direct HiFi assembly; Strainline (X Luo et al., 2022) and HaploDMF (Cai et al., 2022), which have been introduced as viral strain separation methods; and finally iGDA (Z Feng et al., 2021), which can perform both. Software that purposefully collapse similar strains, such as metaMDBG Benoit et al., 2024, were left out of the benchmark.

We tried using all these software on all datasets. Strainline and HaploDMF failed to run in reasonable time on non-viral datasets and were automatically killed after 15 days of processing. Strainline failed to perform strain separation on the HBV-2 dataset within its allowed RAM limit of 50G, probably because of the high coverage. We tried downsampling the dataset but the problem remained.

The reference-based virus phasing tools (haploDMF, iGDA, HairSplitter) were run with the same reference genome as in Cai et al., 2022, namely MT622522.1 for Hepatitis B and MW661279.1 for norovirus.

### Benchmarking evaluation

#### Bacterial haplotypes

The benchmark results on the Zymo-GMS and *V. fluvialis* datasets are summarized in Figure 3 and detailed in Supplementary Table 1. HairSplitter separated better the conspecific strains compared to the original metaFlye assemblies, delivering more comprehensive and accurate assemblies than Strainberry and iGDA. Particularly with Nanopore data, HairSplitter produced the most complete assemblies.

**Figure 3.**
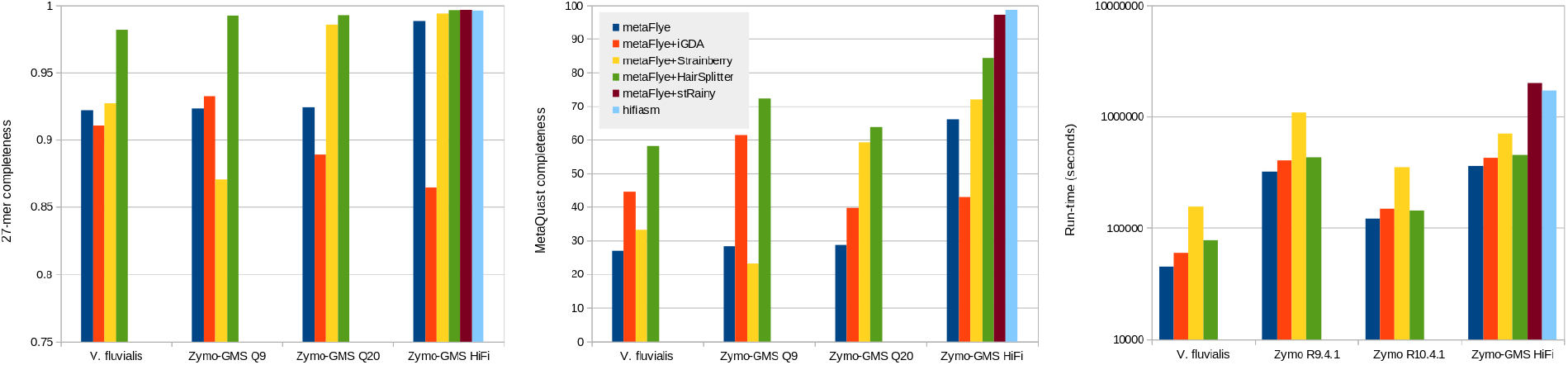
27-mer completeness, MetaQUAST completeness and runtime of different software on the *Vagococcus* and the three Zymo-GMS dataset. The runtimes are the runtimes of the full assembly pipeline (assembly+strain separation) and are represented in log scale.

On HiFi reads, the stRainy, hifiasm and HairSplitter assemblies all had a high k-mer completeness (>98%). However, they showed either a high duplication ratio (for stRainy and hifiasm) or low metaQuast completeness (for HairSplitter) because none managed to duplicate repeated genomic regions to their correct multiplicities (see Supplementary Table 1). This effect was also observed in several Nanopore assemblies, where 27-mer completeness remains high while MetaQUAST completeness is notably lower. Typically, the three almost identical *V. fluvialis* strains were collapsed into one.

The completeness of assemblies in the simulated benchmark is presented in Figure 4, with a detailed evaluation in Supplementary Table 2. The evaluation of iGDA is not depicted because iGDA inexplicably decreased the completeness of the original metaFlye assemblies. Simulations indicated that HairSplitter significantly outperformed Strainberry, particularly in scenarios involving a high number of strains in the metagenome or highly similar strains. The relatively high completeness of the 8-strain Strainberry assembly could be attributed to its high duplication ratio. The completeness of HairSplitter assemblies decreased with the depth of coverage, especially below 20x coverage. The completeness also decreased slightly when the divergence of the strains decreased, though the metaQuast completeness remained high (84%) when assembling two strains with 0.07% divergence. Interestingly, the decline in MetaQUAST completeness when coverage and divergence decreased was more pronounced than the decline in 27-mer completeness, highlighting HairSplitter’s effectiveness in separating divergent regions and its difficulties in duplicating identical regions. This corresponds to the results observed in the Zymo-GMS datasets, where many pairwise divergences of strains were <1%.

**Figure 4.**
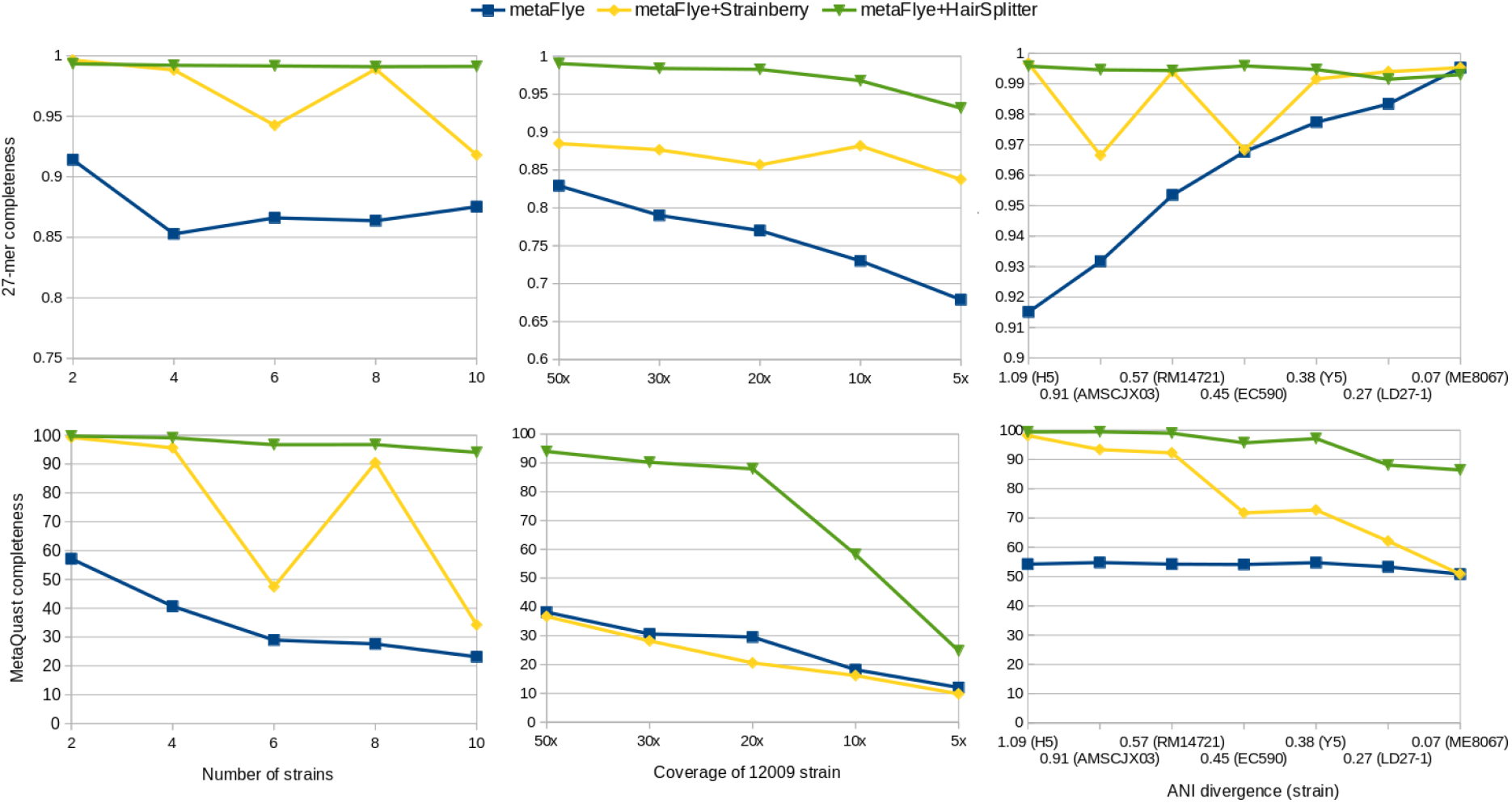
MetaQUAST completeness of assemblies of simulated metagenomes of *E. coli*. On the left, mix of 2 to 10 strains sequenced with 50x coverage were assembled. In the middle, strain 12009 was downsampled in the 10-strains metagenome and completeness of the 12009 strain is measured. On the right, reads of strains of decreasing divergence were mixed with K-12 reads and assembled.

The contigs produced by HairSplitter were found to have a lower number of indels and mismatches compared to iGDA and Strainberry (Sup. Tables 2 and 3). This can be explained by the fact that the groups of reads used by HairSplitter to build the contigs were more homogenous in terms of haplotypes and thus easier to polish. However, all tools produced a significant number of misassemblies when reconstructing a high number of strains. In the case of HairSplitter, these misassemblies were primarily caused by the fact that a few small structural variations were not detected during the graph completion step. In terms of contiguity, all assemblers produced comparable results, although HairSplitter appeared to make slightly more conservative choices than Strainberry, resulting in a slight decrease in contiguity but a lower number of misassemblies (Sup. Table 2 and 3).

#### Viral haplotypes

The completeness results of the benchmark on the viral datasets are depicted Figure 5 and more complete evaluation of assemblies are available in Supplementary Table 3.

**Figure 5.**
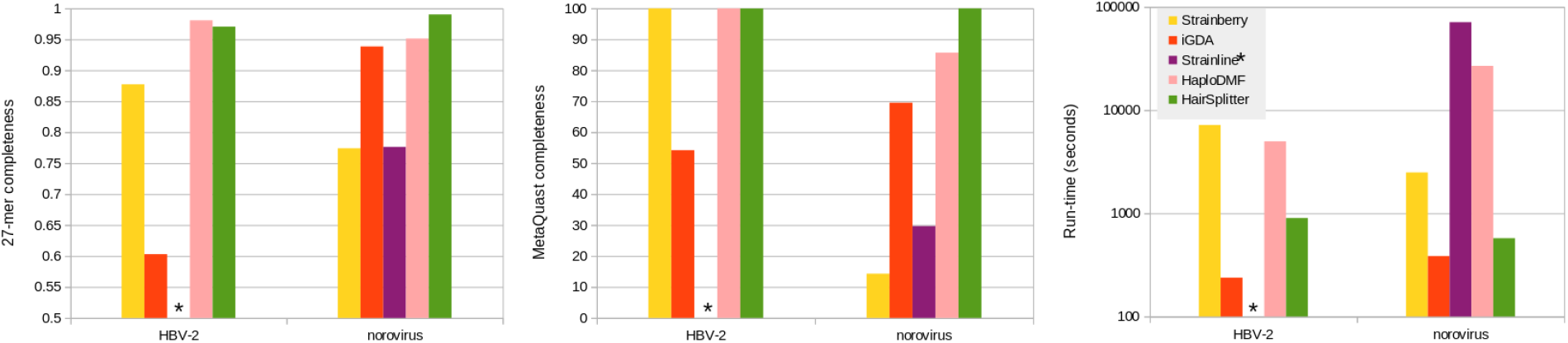
27-mer completeness, MetaQUAST completeness and runtime of different software on the two viral datasets. Note that the runtime is shown in log scale. The Strainline assembly of HBV-2 is not shown because Strainline could not finish on this dataset.

HaploDMF and HairSplitter managed to separate completely the HBV strains according to MetaQUAST. iGDA failed to recover the strains, while Strainberry outputted four different haplotypes instead of two (see supplementary Table 3). We checked that HaploDMF and HairSplitter separated the reads adequately, thus the slight differences in 27-mers completeness stem from polishing errors.

HairSplitter stood out as the sole software capable of successfully recovering all seven strains in the norovirus mix, even capturing the least abundant strain comprising only 1% of the mix. To assess the sensitivity limits of HairSplitter in the viral context, we conducted two additional experiments within the norovirus mix. In the first experiment, we decreased the relative abundance of the rarest strain to 0.1%, while maintaining 50x coverage by uniformly increasing the coverage of the other strains. Remarkably, HairSplitter still achieved complete recovery (99.99% MetaQUAST completeness) of the rarest strain. The limited amount of data prevented us to further reduce the strain’s relative abundance. In the second experiment, we uniformly diminished the coverage of all strains. The rarest strain was entirely recovered (99.99% MetaQUAST completeness) when covered at *≥*40x, only the most divergent part of the virus was recovered (26.4% MetaQUAST completeness) at coverage 20x and 30x, and the strain was not recovered at all at 10x coverage. The primary determinant of HairSplitter’s sensitivity thus seems to be absolute coverage rather than the strain’s relative coverage.

## Discussion

In this manuscript, we introduce HairSplitter, a pipeline to assemble haplotypes separately using an input assembly and long reads. The pipeline includes two main novelties, a program that completes an assembly graph and a read separation procedure. HairSplitter proved useful when dealing with noisy data (*≥* 1% error rate), whereas its usefulness on HiFi reads compared to specialised programs such as hifiasm or stRainy is debatable. We show that HairSplitter can effectively separate several highly similar strains in both bacterial and viral contexts. Compared to the state of the art, HairSplitter can deal with a higher number of strains, lower relative abundances and lower strain divergence, while maintaining a low computational cost.

HairSplitter encounters a major limitation when strains have many identical regions. In these regions, it is not possible to assign reads to specific haplotype groups, making it necessary to duplicate the homozygous regions to their correct multiplicity in order to fully recover the strains. This study demonstrates that this is a challenging problem that current assemblers are not able to successfully address in the HiFi dataset. Further investigation will be needed to solve this issue. A lead could be to use astutely the topology of the assembly graph.

A direction for future work would also be to generalize the assembly graph completion module. The idea of the module is to make sure all reads align end-to-end onto the assembly graph. We believe such a module could be useful to improve many assemblies. However, the version implemented for now in HairSplitter is very basic and does not perform well in repeated, complicated regions of the graph. A more sophisticated module could involve local reassembly and iterative graph completion. Such work has started and can be followed on GitHub: github.com/rolandfaure/genometailor

Since HairSplitter is already successful at separating both bacterial and viral haplotypes, we expect to be able to extend this work naturally towards the phasing of polyploid organisms, motivated by the fact that for now, polyploid genome assembly requires highly precise illumina or HiFi reads (Kong et al., 2023). For this particular case, some extra information could be leveraged to improve the HairSplitter pipeline, such as the fact that all haplotypes are expected to be equally abundant and that the total number of haplotype is usually known.

## Reproducibility and data availablility

The HairSplitter code can be found on GitHub at https://github.com/rolandfaure/hairsplitter.

The experiments were run with Flye 2.9.2-b1786, hifiasm HairSplitter v1.9.4, HaploDMF commit a07d082c3, Strainline commit 8d26341, iGDA commit 54ecec9, Strainberry v1.1, stRainy commit 34573cd, hifiasm-meta v0.3-r063.2, minimap2 v2.26-r1175 and Quast v5.2.0.

HBV sequencing reads can be found under accession number ERR3253560 in SRA. The seven norovirus sets of reads can be found under accession numbers SRR13951181, SRR13951181, SRR13951186, SRR13951185, SRR13951184, SRR13951165 and SRR13951160. The *Vagococcus fluvialis* data are accessible under project PRJNA755170 in SRA. The Zymo-GMS sequencing data can be found under accession numbers SRR17913200, SRR17913199 and SRR13128013.

All the assemblies, simulated data and command lines used are available on Zenodo, DOI 10.5281/zenodo.10495033, https://zenodo.org/records/11639887.

## Acknowledgments

We thank Ulysse Faure for his mathematical help. Alexandros Vasilikopoulos, Andrew Woodruff and Alessandro Derzelle tested HairSplitter and kindly helped debugging.

We acknowledge the GenOuest bioinformatics core facility (https://www.genouest.org) for providing the computing infrastructure. The programs Tablet (Milne et al., 2013) and Bandage (RR Wick et al., 2015) were used to visualize data while developing HairSplitter.

For the purpose of Open Access, a CC-BY public copyright licence has been applied by the authors to the present document and will be applied to all subsequent versions up to the Author Accepted Manuscript arising from this submission

