## Supplementary Tables for "HairSplitter: haplotype assembly from long, noisy reads"

Peer Community In
Mathematical & Computational Biology

RESEARCH ARTICLE

Open Access
Open Peer-Review
Open Data
Open Code

### Supplementary material - HairSplitter: haplotype assembly from long, noisy reads

Roland Faure [1,2], Dominique Lavenier[1] & Jean-François Flot[2,3]

[1] Univ. Rennes, INRIA RBA, CNRS UMR 6074, Rennes, France
[2] Service Evolution Biologique et Ecologie, Université libre de Bruxelles (ULB), Brussels, Belgium
[3] Interuniversity Institute of Bioinformatics in Brussels – (IB)[2], Brussels, Belgium

**Cite as:**
xxx

**Correspondence:**

**Recommender:**
FirstName FamilyName

**Reviewers:**
FirstName FamilyName and two anonymous reviewers

This version of the article has not yet been peer-reviewed by *Peer Community In Mathematical and Computational Biology* (https://doi.org/xxx/xxx)

#### Abstract

**Motivation:** Long-read assemblers face challenges in discerning closely related viral or bacterial strains, often collapsing similar strains in a single sequence. This limitation has been hampering metagenome analysis, where diverse strains may harbor crucial functional distinctions.

**Results:** We introduce a novel software, HairSplitter, designed to retrieve strains from a strain-oblivious assembly and long reads. The method uses a custom variant calling process to operate with erroneous long reads and introduces a new read binning algorithm to recover an a priori unknown number of strains. On noisy long reads, HairSplitter can recover more strains while being faster than state-of-the-art tools, both in the viral and the bacterial case.

**Availability:** HairSplitter is freely available on GitHub at github.com/RolandFaure/HairSplitter.

**Contact:**

***Keywords:*** Metagenomes; Metaviromes; Haplotyping; Genome assembly; Strain separation

| | | Complet--eness (%) | Duplication ratio | NGA50 | #misas--semblies | #mismatches per 100 kbp | # indels per 100 kbp | Assembly length (Mb) |
|---|---|---|---|---|---|---|---|---|
| *Vagococcus fluvialis* | metaFlye | 26.90 | 1.097 | - | 40 | 340.14 | 383.57 | 4.4 |
| | metaFlye + iGDA | 44.530 | 1.151 | 1313 | 2 | 115.57 | 391.70 | 7.4 |
| | metaFlye + Strainberry | 33.112 | 1.119 | - | 45 | 77.71 | 510.25 | 5.5 |
| | metaFlye + HairSplitter | 58.150 | 1.066 | 19881 | 31 | 102.49 | 410.95 | 9.1 |
| Zymo-GMS Q9* | metaFlye | 28.365 | 1.048 | - | 66 | 286.86 | 38.80 | 7.4 |
| | metaFlye + iGDA | 61.293 | 1.489 | 12450 | 21 | 225.67 | 41.30 | 22.6 |
| | metaFlye + Strainberry | 23.166 | 1.072 | - | 45 | 191.51 | 51.65 | 6.1 |
| | metaFlye + HairSplitter | 72.293 | 1.105 | 9974 | 24 | 65.77 | 39.60 | 19.8 |
| Zymo-GMS Q20* | metaFlye | 28.742 | 1.051 | - | 62 | 300.25 | 34.41 | 7.5 |
| | metaFlye + iGDA | 39.650 | 1.118 | - | 8 | 181.55 | 28.23 | 11.0 |
| | metaFlye + Strainberry | 59.197 | 1.138 | 28421 | 66 | 193.55 | 42.42 | 16.7 |
| | metaFlye + HairSplitter | 63.837 | 1.022 | 12000 | 43 | 40.01 | 21.69 | 16.2 |
| Zymo-GMS HiFi* | metaFlye | 66.064 | 1.076 | 79832 | 39 | 92.55 | 6.65 | 17.6 |
| | metaFlye + iGDA | 42.996 | 1.515 | 17104 | 15 | 102.70 | 9.96 | 16.1 |
| | metaFlye + Strainberry | 72.016 | 1.142 | 53249 | 46 | 57.53 | 6.92 | 20.4 |
| | metaFlye + HairSplitter | 84.418 | 1.286 | 25851 | 69 | 32.35 | 12.16 | 26.9 |
| | metaFlye + stRainy | 97.078 | 1.737 | 41195 | 47 | 44.15 | 12.26 | 41.8 |
| | hifiasm | 98.732 | 1.911 | 288422 | 82 | 30.07 | 4.99 | 46.7 |

**Table 1.** metaQuast metrics of the bacterial assemblies obtained from experimental data. *metrics are computed with respect to the 5 *E. coli* strains, not the complete dataset - the assembly length is the aligned assembly length on the *E. coli* reference

| | | Complet- -eness (%) | Duplication ratio | NGA50 | #misas- -semblies | #mismatches per 100 kbp | # indels per 100 kbp | Assembly length (Mb) |
|---|---|---|---|---|---|---|---|---|
| # strains | | | | | | | | |
| 2 | metaFlye | 57.137 | 1.043 | 61559 | 22 | 216.18 | 216.53 | 6.1 |
| | metaFlye + Strainberry | 99.268 | 1.074 | 701492 | 11 | 24.83 | 64.73 | 10.8 |
| | metaFlye + HairSplitter | 99.716 | 1.008 | 396746 | 1 | 8.21 | 42.73 | 10.2 |
| 4 | metaFlye | 40.666 | 1.071 | - | 50 | 562.91 | 268.83 | 8.9 |
| | metaFlye + Strainberry | 95.631 | 1.148 | 251144 | 39 | 93.07 | 81.88 | 22.3 |
| | metaFlye + HairSplitter | 99.109 | 1.064 | 109320 | 39 | 25.74 | 50.83 | 21.3 |
| 6 | metaFlye | 28.949 | 1.087 | - | 63 | 585.20 | 264.89 | 9.4 |
| | metaFlye + Strainberry | 47.430 | 1.108 | 8151 | 90 | 289.19 | 92.23 | 15.7 |
| | metaFlye + HairSplitter | 96.717 | 1.086 | 77500 | 125 | 52.55 | 56.23 | 31.1 |
| 8 | metaFlye | 27.599 | 1.051 | - | 76 | 527.44 | 277.85 | 11.5 |
| | metaFlye + Strainberry | 90.438 | 1.533 | 83755 | 157 | 179.31 | 172.25 | 54.7 |
| | metaFlye + HairSplitter | 96.759 | 1.180 | 45253 | 244 | 87.74 | 84.97 | 45.1 |
| 10 | metaFlye | 23.130 | 1.036 | - | 79 | 469.27 | 277.71 | 11.7 |
| | metaFlye + Strainberry | 34.207 | 1.095 | - | 175 | 363.06 | 137.01 | 18.2 |
| | metaFlye + HairSplitter | 94.045 | 1.192 | 41223 | 262 | 93.74 | 66.47 | 54.5 |
| coverage | | | | | | | | |
| 30x* | metaFlye | 30.618 | 1.032 | - | 1 | 719.80 | 55.27 | 1.7 |
| | metaFlye + Strainberry | 28.188 | 1.085 | - | 2 | 347.21 | 127.06 | 1.7 |
| | metaFlye + HairSplitter | 90.206 | 1.170 | 40243 | 19 | 143.08 | 41.63 | 5.8 |
| 20x* | metaFlye | 29.522 | 1.018 | - | 6 | 859.11 | 76.03 | 1.7 |
| | metaFlye + Strainberry | 20.562 | 1.037 | - | 3 | 274.86 | 102.01 | 1.2 |
| | metaFlye + HairSplitter | 87.948 | 1.093 | 37879 | 22 | 130.95 | 77.82 | 5.2 |
| 10x* | metaFlye | 18.201 | 1.005 | - | 1 | 498.17 | 85.10 | 1.0 |
| | metaFlye + Strainberry | 16.178 | 1.007 | - | 1 | 347.06 | 181.70 | 0.9 |
| | metaFlye + HairSplitter | 58.172 | 1.054 | 10763 | 6 | 214.47 | 117.59 | 3.3 |
| 5x* | metaFlye | 12.020 | 1.010 | - | 2 | 849.88 | 201.29 | 0.6 |
| | metaFlye + Strainberry | 9.807 | 1.013 | - | 2 | 422.61 | 273.49 | 0.5 |
| | metaFlye + HairSplitter | 24.746 | 1.082 | - | 2 | 464.50 | 197.55 | 1.5 |
| divergence | | | | | | | | |
| H5 | metaFlye | 54.246 | 1.002 | 19206 | 22 | 324.97 | 29.23 | 5.1 |
| (1.09%) | metaFlye + Strainberry | 98.157 | 1.001 | 652572 | 2 | 324.97 | 1.35 | 9.3 |
| | metaFlye + HairSplitter | 99.419 | 1.008 | 294365 | 6 | 8.08 | 15.25 | 9.4 |
| AMSCJX03 | metaFlye | 54.783 | 1.001 | 18675 | 17 | 254.48 | 28.42 | 5.0 |
| (0.91%) | metaFlye + Strainberry | 93.390 | 1.003 | 279448 | 3 | 0.73 | 1.98 | 8.6 |
| | metaFlye + HairSplitter | 99.456 | 1.002 | 387661 | 11 | 10.96 | 23.29 | 9.2 |
| RM74721 | metaFlye | 54.254 | 1.000 | 19360 | 19 | 132.60 | 11.73 | 5.0 |
| (0.57%) | metaFlye + Strainberry | 92.256 | 1.006 | 380826 | 1 | 2.97 | 8.57 | 8.6 |
| | metaFlye + HairSplitter | 98.957 | 1.007 | 265482 | 14 | 15.54 | 29.08 | 9.2 |
| EC590 | metaFlye | 54.132 | 1.000 | 17337 | 10 | 117.79 | 12.85 | 5.0 |
| (0.45%) | metaFlye + Strainberry | 71.749 | 1.003 | 156627 | 10 | 9.29 | 1.72 | 6.6 |
| | metaFlye + HairSplitter | 95.697 | 1.024 | 190750 | 6 | 6.46 | 24.51 | 9.0 |
| Y5 | metaFlye | 54.736 | 1.002 | 22104 | 21 | 63.85 | 9.38 | 5.2 |
| (0.38%) | metaFlye + Strainberry | 72.758 | 1.006 | 181387 | 13 | 8.64 | 3.28 | 6.9 |
| | metaFlye + HairSplitter | 97.154 | 1.024 | 253619 | 8 | 40.80 | 52.36 | 9.4 |
| LD27-1 | metaFlye | 53.295 | 1.001 | 19411 | 8 | 43.83 | 5.33 | 5.0 |
| (0.27%) | metaFlye + Strainberry | 62.101 | 1.004 | 112749 | 13 | 7.41 | 3.53 | 5.8 |
| | metaFlye + HairSplitter | 88.101 | 1.055 | 137245 | 2 | 95.36 | 97.47 | 8.6 |
| ME8067 | metaFlye | 50.820 | 1.000 | 47356 | 8 | 10.29 | 3.19 | 4.7 |
| (0.07%) | metaFlye + Strainberry | 50.820 | 1.000 | 47356 | 8 | 10.29 | 3.19 | 4.7 |
| | metaFlye + HairSplitter | 86.419 | 1.064 | 131660 | 0 | 84.75 | 80.83 | 8.5 |

**Table 2.** metaQuast metrics of the bacterial assemblies obtained from simulated Nanopore R10.4.1 data. * The metrics displayed for the downsampled datasets are the metrics computed with respect to the downsampled strain, and not with respect to the complete 10 strains.

| | | Completeness (%) | Duplication ratio | NGA50 | #misassemblies | # mismatches per 100 kbp | # indels per 100 kbp |
|---|---|---|---|---|---|---|---|
| HBV-2 | Strainberry | 99.984 | 2.174 | 4504 | 3 | 881.59 | 1562.50 |
| | iGDA | 54.174 | 1.001 | 1081 | 0 | 201.15 | 229.89 |
| | Strainline | | | | | | |
| | HaploDMF | 99.984 | 1.000 | 3207 | 0 | 15.58 | 93.46 |
| | HairSplitter | 99.953 | 1.001 | 3209 | 0 | 46.72 | 109.02 |
| norovirus | Strainberry | 14.283 | 1.000 | - | 0 | 52.97 | 13.24 |
| | iGDA | 69.514 | 1.548 | 2838 | 0 | 112.55 | 15.83 |
| | Strainline | 29.659 | 5.787 | 7541 | 0 | 479.44 | 136.67 |
| | HaploDMF | 85.702 | 1.000 | 7549 | 0 | 165.60 | 26.50 |
| | HairSplitter | 100.000 | 1.038 | 7550 | 0 | 107.57 | 35.96 |

**Table 3.** metaQuast metrics of the viral assemblies. The Strainline metrics are empty for the HBV-2 dataset because Strainline crashed.

| Number of strains | | metaFlye assembly | metaFlye assembly after graph completion |
|---|---|---|---|
| 2 | N50 | 374204 | 374204 |
| | #misassemblies | 22 | 20 |
| 4 | N50 | 136064 | 47179 |
| | #misassemblies | 50 | 17 |
| 6 | N50 | 78959 | 36209 |
| | #misassemblies | 63 | 12 |
| 8 | N50 | 73412 | 32985 |
| | #misassemblies | 76 | 18 |
| 10 | N50 | 55801 | 26906 |
| | #misassemblies | 79 | 19 |

**Table 4.** N50 and metaQuast-measured number of misassemblies of simulated datasets with varying number of *E. coli* strains, before and after completing the assembly graph. Since the completion step breaks contigs, the N50 diminishes. The number of misassemblies diminishes with graph completion.
